# The return of the trophic chain: fundamental vs realized interactions in a simple arthropod food web

**DOI:** 10.1101/324178

**Authors:** Inmaculada Torres-Campos, Sara Magalhães, Jordi Moya-Laraño, Marta Montserrat

## Abstract

The mathematical theory describing small assemblages of interacting species (community modules or motifs) has proved to be essential in understanding the emergent properties of ecological communities. These models use differential equations to study pairwise interactions between species. However, as community modules become more complex, it is not certain that all potential interactions will be effectively realized. Here, we use community modules to experimentally explore whether the number of trophic links among species scales with community complexity (i.e., by adding species known to feed on each other from pair-wise trials). To investigate this question, we used a simple mite community present in avocado orchards (*Persea americana*), composed of two predators (*Euseius stipulatus* and *Neoseiulus californicus*), one herbivore as shared prey (*Oligonychus perseae*), and pollen of *Carpobrotus edulis* as an alternative food resource. These configurations enabled the potential for (intraguild) predation and (apparent) competition to be expressed. Using a series of controls, we assessed whether the presence of one species affected the survival of another, or its conversion of food into offspring). We found that increasing the number of potential interactions did not result in more complex realized community modules. Instead, all communities were reduced to one or two linear trophic chains. Our results show that trophic links assumed to occur when species are assembled in pairs do not necessarily occur when other components of the community are present. Consequently, food-web structure may be unrealistic in theoretical community modules that are parameterized based on pair-wise interactions observed when alternative prey is absent. This further suggests the need for empirical research to work in concert with theoretical approaches to develop more realistic and predictive food-web models.

## Introduction

Community ecology initially conceptualized trophic interactions as linear chains (Elton 1927), with an upper level potentially controlling the densities of the level immediately below, thus generating a trophic cascade (Hairston *et al.* 1960; Oksanen *et al.* 1981; Carpenter *et al.* 1985). However, it is now accepted that most communities do not follow this pattern as organisms are imbedded in complex food webs, blurring the notion of a trophic guild (*sensu* trophic coherence, Johnson et al. 2014) and the notion that widespread omnivory destabilize food webs (Polis & Holt 1992; Polis & Strong 1996).

Food webs can be decomposed into “community modules” (i.e. “small number of species (e.g. three to six) linked in a specified structure of interactions, Holt 1997). Community modules are similar to motifs, used in network studies (e.g., Bascompte & Melián 2005, Prill et al 2005). Among those, intraguild predation (IGP), in which two consumers (the intraguild predator and the intraguild prey, hereafter IG-predator and IG-prey) not only compete for a shared resource but also engage in predator-prey interactions (Polis *et al.* 1989), and apparent competition, in which two non-competing prey share a common predator (Holt 1977; 1997), are the most common (Bascompte & Melián 2005).

Whether and how often species engage in intraguild predation or apparent competition strongly affects the long-term persistence (i.e. “temporal stability in community composition”, Pimm 1984) of communities. Theory predicts that intraguild predation destabilizes communities because it reduces the parameter space where coexistence of IG-predator, IG-prey and shared prey is possible (Holt & Polis 1997), compared to that of trophic chain models (Oksanen *et al.* 1981). In most models with intraguild predation the possibility of 3-species persistence requires the IG-prey to be superior to IG-predators at exploitation competition for the shared resource (Holt 1997; Mylius *et al.* 2001; Kondoh 2008, among others). Furthermore, the occurrence of intraguild predation leads to a less efficient control of the shared prey populations because the IG-prey’s equilibrium abundance is expected to decline towards extinction with increasing productivity (Mylius *et al.* 2001). This is especially important in agricultural systems, inherently highly productive, in which the introduction of several biocontrol agents may in fact reduce pest control. Curiously, empirical studies, mostly stemming from such systems, show that variable effects of intraguild predation on populations of the shared prey (e.g., Rosenheim *et al.* 1995; Janssen *et al.* 2006; Vance-Chalcraft *et al.* 2007; Messelink & Janssen 2014).

Some factors may, however, reduce such instability by promoting species coexistence, which generally occurs when conditions under which predator-predator interactions occur are constrained (reviewed in Novak 2013, appendix S1). For example, 3-species coexistence is enhanced if predators engage in IGP only when competition for the shared prey is high (Křivan 2000), or the shared prey is less profitable than the intraguild prey (Křivan & Diehl 2005). Other studies suggest that the inclusion of habitat structure (Janssen *et al.* 2007), inducible defences (Kratina *et al.* 2010; Nakazawa *et al.* 2010) or temporal refuges (Amarasekare 2008) increases the persistence of IGP communities, although this may depend on which species use refuges (Liu & Zhang 2013). Moreover, stage structure in the intraguild prey promotes 3-species coexistence, either by providing a stage refuge (Mylius *et al.* 2001; Rudolf & Armstrong 2008) or by inducing ontogenetic niche shifts in the predator (Hin *et al.* 2011). However, in all cases, the models still predict that overall community persistence is lower than that of a simple trophic chain. This lack of temporal stability is corroborated by empirical laboratory studies (Diehl & Feißel 2000; Montserrat *et al.* 2008b), but runs counter the ubiquity of intraguild predation and trophic level omnivory in natural systems (Bascompte & Melián 2005; Gagnon *et al.* 2011).

Discrepancies between IGP theory and empirical data suggest that some assumptions of theoretical models are not met in natural systems. In an effort to bring IGP models closer to real systems, while maintaining mathematical tractability, researchers have tested how the incorporation of an alternative food source affects the persistence of IGP communities (Heithaus 2001; Daugherty *et al.* 2007; Holt & Huxel 2007; Rudolf 2007). The general prediction is that providing alternative food to the intraguild prey leads to wider parameter regions of species coexistence (Daugherty *et al.* 2007; Holt & Huxel 2007), even if competitive superiority of IG-prey is precluded (Faria & Costa 2010). Instead, alternative food for the intraguild predator destabilizes the community (Daugherty *et al.* 2007; Holt & Huxel 2007). However, in the latter case, if the alternative food quality is high, then the intraguild predator may switch to feeding on the alternative resource, whereas the intraguild prey feeds on the shared prey (Ibid.). This again promotes coexistence by bringing the community structure closer to two linear food chains. Thus, a prevailing outcome of the ecological theory is that domains for persistence of communities with IGP increase when the strength of intraguild predation decreases. Indeed, weak interactions have long been recognized to stabilize ecosystems by dampening oscillations between consumers and resources, thereby decreasing the probability of species extinctions (McCann *et al.* 1998), and thus promoting community persistence (May 1972; Pimm & Lawton 1978; Paine 1992; McCann *et al.* 1998; Emmerson & Yearsley 2004; Neutel *et al.* 2007; Gellner & McCann 2012; 2016). However, it remains unclear how the addition of species into a community alters all other trophic interactions in the network (e.g., Bascompte et al 2006).

To address this question, we have considered if the fundamental trophic niches of species (i.e., with all their potential interactions; Elton 1927) are always realized (Hutchinson 1957). Specifically, we explore how pairwise trophic interactions between species are modified by the inclusion of other species in a simple community. We focus on *predation rate* (in here, number of individuals consumed per day) as it is an excellent proxy for trophic interaction strength, and is used both in ecological modelling (e.g. the equivalent to the “catching efficiencies” in Kuijper *et al.* 2003) and in empirical research (Wootton & Emmerson 2005; Novak & Wootton 2010; Novak 2013). Measurements of other relevant non-trophic interactions, such as *competition*, would require experiments at the population and community level that are beyond the scope of this manuscript.

Our core hypothesis was that increasing the number of species that are known to interact when no alternative food is available will increase the number of realized links in the more complex community (Box 1A). We mimicked different community modules (Sensu Holt 1997) of increasing complexity using a community composed of two predatory mite species as intraguild predators (*Euseius stipulatus* and *Neoseiulus californicus*, Acari: Phytoseiidae), one species of herbivore mite as the shared prey (*Oligonychus perseae*, Acari: Tetranychidae), and pollen (of several anemophilous species) as alternative food (González-Fernández *et al.* 2009), all of which occur in the leaves of crops of avocado plants (*Persea americana*) in Southestern Spain. Previous pairwise experimental designs have shown that the interaction between *N. californicus* and *O. perseae* is stronger (i.e., predation rates are higher) than that between *E. stipulatus* and this same prey (González-Fernández et al. 2009). Moreover, pollen is an optimal food source for *E. stipulatus* but not for *N. californicus* (Ferragut et al. 1987; González-Fernández et al. 2009). Finally, *E. stipulatus* and *N. californicus* engage in size-dependent predator-prey interactions (Abad-Moyano *et al.* 2010). This knowledge was used to build predictions on realized trophic links occurring in this system across community modules of increasing complexity (Box 1B). Specifically, we predicted that: *i*) in “trophic chain” community configurations, both predator species will interact with the herbivore (Box 1B, a.1.1. and a.1.2.); *ii*) in “apparent competition” community configurations, only *E. stipulatus* will interact with both the herbivore and pollen (Box 1B, b.1.1. and b.1.2.); *iii*) in “intraguild predation” community configurations, both IG-predator species will interact with the IG-prey and the herbivore (Box 1B, c.1.1. and c.1.2.); and iv) in “Intraguild predation and apparent competition” community configurations, only adults and juveniles of *E. stipulatus* will establish trophic links with pollen (Box 1B, d.1.1. and d.1.2.). These predictions were then tested through a series of experimental treatments to assess which interactions were realized within each community module, by measuring IG-prey/herbivore mortality and how consumption of prey translates into predator fecundity as a result of these interactions. Specifically, we examined a) whether (IG-)predators feed on each prey type; b) whether predation of (IG-)predators on one prey type is affected by the presence of the other; c) whether predation of (IG-)predators on both prey, and of IG-prey on the herbivore, is affected by the presence of alternative food; d) whether the presence of alternative food affects predation of (IG-)predators on the two types of prey when they are together; e) number of eggs produced by (IG-)predators when feeding on each prey type; and f) whether egg-production is additive when (IG-)predators have more than one food type available.

## Material and Methods

All cultures and experiments were done in a climate chamber at 25±1°C, 65±5% RH and 16:8h L:D (Light:Dark).

### Mite cultures

Cultures of the predatory mite *E. stipulatus* were started in 2007 from ca. 300 individuals collected from avocado trees located in the experimental station of “La Mayora”. Rearing units consisted of three bean plants (*Phaseolus vulgaris* L.) with 6-10 leaves, positioned vertically, with the stems in contact with sponges (*ca.* 30 × 20 × 5 cm) covered with cotton wool and a plastic sheet (27 × 17 cm), and placed inside water-containing trays (8 L, 42.5 × 26 × 7.5 cm). The plant roots were in contact with the water, and the aerial parts were touching each other, forming a tent-like three-dimensional structure, where individuals could easily walk from one plant to the other. Cotton threads were placed on the leaves, to serve as oviposition sites for the females. Mites were fed *ad libitum* twice a week with pollen of *Carpobrotus edulis* (cat´s claw) spread on leaves with a fine brush. *Euseius stipulatus* is able to develop and reproduce on this food source (Ferragut *et al.* 1987). Every three weeks, new rearings were made by transferring, leaves with mites and the cotton threads filled with eggs to a new unit. The culture was found to be contaminated a few times with *Tyrophagus* spp., a detritivorous mite species. In such instances, instead of moving entire leaves, adult *E. stipulatus* females (ca. 300) were collected individually and transferred to the new rearing unit.

The *N. californicus* population was obtained from Koppert Biological Systems S.L. in bottles of 1000 individuals (Spical^®^). Colonies were kept on detached bean leaves infested with *Tetranychus urticae* that were placed on top of inverted flower-pots (20 cm Ø) inside water-containing trays.

The herbivore *Oligonychus perseae* was not maintained in a laboratory culture due to technical difficulties in preserving detached avocado leaves. They were thus collected from the field on a regular basis from avocado orchards located in the experimental station of “La Mayora”.

Pollen of *C. edulis* was obtained from flowers collected in the experimental station. Stamens dried in a stove at 37°C for 48h, then sieved (350 µm).

### Community modules

Experimental arenas to test the outcome of community modules have been described in detail in Guzmán *et al.* (2016a). Briefly, a hole (6.5 cm Ø) was cut in a petri dish (9 cm Ø), turned upside down, and then filled with an avocado leaf disc (7.5 cm Ø). The borders were glued to a clay ring. Inside the petri dish, wet cotton wool ensured enough humidity to keep leaves turgid. Petri dishes were then sealed with parafilm^®^. To prevent individuals from escaping, a ring of Tanglefoot^®^ was applied along the outer margin of the leaf disc. Cohorts of *E. stipulatus* were made by transferring with a fine brush 400 eggs from the rearings to 2-3 bean leaves placed on top of sponges (30 × 20 × 5 cm, approx.) covered with cotton wool, inside water-containing trays (3.5 L), and with pollen of *C. edulis* as food. Cohorts of *N. californicus* were made by placing 100 females during 48 h on 2-3 bean leaves infested with *Tetranychus urticae* in containers similar to those used for the cultures. 10-14 days after egg hatching, gravid predator females were randomly taken from these cohorts, and starved for 16 h in experimental containers similar to those above. Starvation was done to standardize hunger among individuals, and to ensure that egg production in tested females was not obtained from food ingested prior to the experiment. Predator juveniles (2-3 days old since hatching) were taken from the cohorts when needed. Arenas containing the herbivore were done as follows: Ten females of *O. perseae* were let to build nests and lay eggs on experimental arenas during 4 days. The number of nests and eggs per nest on each arena was counted at the onset of the experiment. Pollen in arenas assigned to treatments with alternative food was supplied *ad libitum*, using a fine brush.

We performed experiments using two ‘community blocks’, depending on the identity of the top predator (*N. californicus* or *E. stipulatus*). Throughout the text, the identity of (IG)-predator and (IG)-prey will be indicated using the subscripts “ES” for *E. stipulatus* and “NC” for *N. californicus*. Increased complexity in each of the two community blocks was mimicked through the combination of the presence / absence of 4 factors: predator/IG-predator, IG-prey, herbivore and alternative food. This resulted in the community modules (Sensu Holt 1997) depicted in the X-axis of figures 1 and 2. These modules were: 1. <u>Trophic</u> <u>chain</u>: either one *E. stipulatus* or *N. californicus* female was introduced in arenas containing 10 females of *O. perseae* (treatment # 1 in Figs 1 and 2). Arenas containing either one *E. stipulatus* or one *N. californicus* female without herbivores (treatment # 2), and containing 10 *O. perseae* females without predators (treatment # 3) were done as controls for predator oviposition rate and prey natural mortality, respectively. 2. <u>Apparent competition</u>: arenas consisted of one female of either *E. stipulatus* or *N. californicus*, 10 females of *O. perseae*, and pollen of *C. edulis* supplied *ad libitum* (treatment # 4). Similar arenas but without the herbivores (treatment # 5) were made as controls for oviposition rates of predators on pollen only, and without the IG-predator (treatment # 6) to assess potential effects of pollen on the survival of the herbivore. 3. <u>Intraguild predation</u>: Because IGP is usually associated with size differences between contestants, IG-predators and IG-prey consisted of adult females and heterospecific juveniles, respectively. Arenas consisted of 10 *O. perseae* females, either one *E. stipulatus* or *N. californicus* female, acting as the IG-predators, and 10 heterospecific juveniles, acting as the IG-prey (treatment # 7). Additionally, control treatments were done to evaluate: the predation/mortality rate of *O. perseae* in the presence of IG-prey but not of IG-predator (treatment # 8); the mortality rate of IG-prey in the absence of both IG-predator and prey (treatment # 9), and in the presence of IG-predator but not of herbivores (treatment # 10). 4. <u>Intraguild predation - Apparent competition</u>: Arenas consisted of 10 *O. perseae* females, either one *E. stipulatus* or *N. californicus* female, acting as the IG-predators, 10 heterospecific juveniles, acting as the IG-prey, and pollen of *C. edulis* as alternative food, supplied *ad libitum* (treatment # 11). Similar arenas to those above but i) without IG-predators (treatment # 12), ii) without herbivores (treatment # 13), and iii) without IG-predators and herbivores (treatment # 14), were done to evaluate predation of IG-prey on the herbivore in the presence of pollen, predation of IG-predators on IG-prey in the presence of pollen, and mortality of IG-prey in the presence of pollen, respectively.

**Figure 1.**
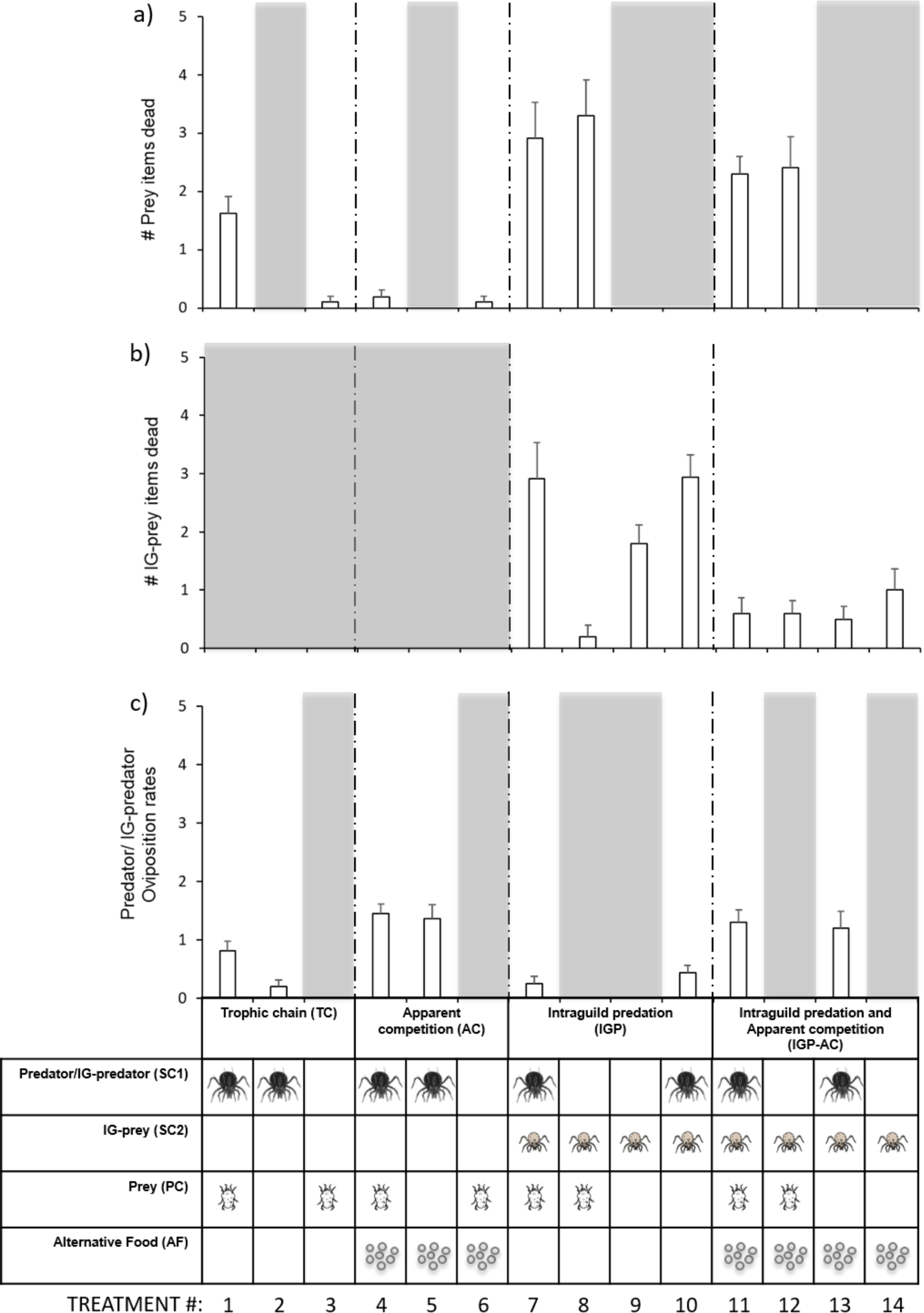
Mortality rates (average ± S.E.) of a) herbivore prey (*Oligonychus perseae* females) and b) IG-prey (*Neoseiulus californicus* juveniles), and c) oviposition rates (average ± S.E.) of IG-predators (*Euseius stipulatus* females), in 14 different treatments defined by presence or absence of either IG-predators, IG-prey, herbivores or alternative food (pollen), depicted in the lower part of the figure, that mimicked four different community configurations and their respective controls.

**Figure 2.**
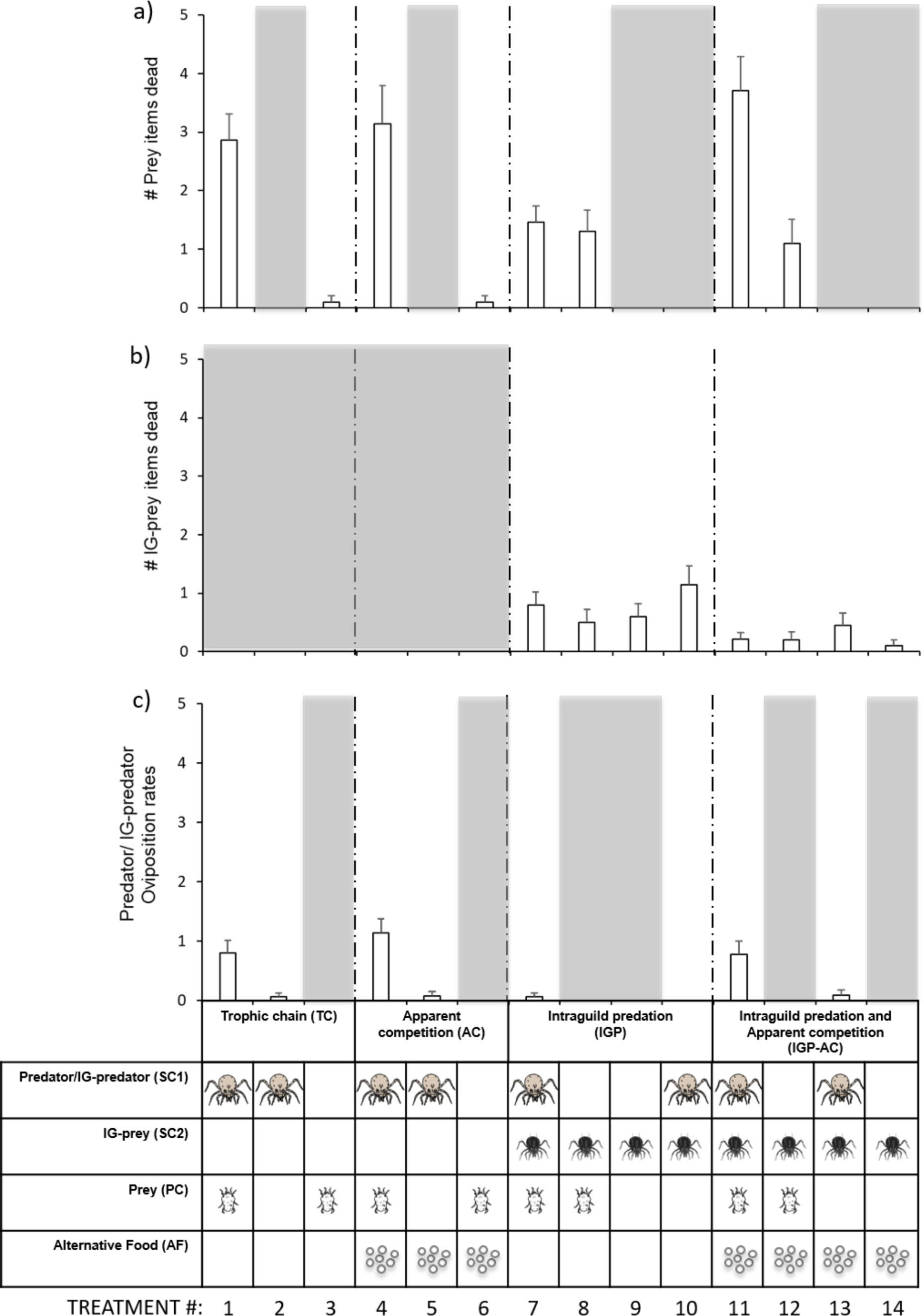
Mortality rates (average ± S.E.) of a) herbivore prey (*Oligonychus perseae* females) and b) IG-prey (*Euseius stipulatus* juveniles), and c) oviposition rates (average ± S.E.) of IG-predators (*Neoseiulus califonicus* females), in 14 different treatments defined by presence or absence of either IG-predators, IG-prey, herbivores or alternative food (pollen), depicted in the lower part of the figure, that mimicked four different community configurations and their respective controls.

Twenty-four hours later, the number of dead herbivores/IG-prey (predation/mortality rate), and the number of eggs laid by predators/IG-predators (oviposition rate) were recorded. Each treatment was replicated between 10 to 18 times.

### Data analyses

Analyses were done separately for communities where either *E. stipulatus* or *N. californicus* acted as the top-predator. Predation rates on herbivores and on IG-prey, and rates of oviposition of IG-predators, were analysed using Generalized Lineal Models (GLM) assuming a Poisson distribution as the distribution of data is expected to be skewed towards low rather than high numbers, and a Log-link function as no overdispersion of the data was detected. All the analyses were 3 full-factorial designs; the main factors that were included in each analysis are explained below. We followed a backward elimination procedure as follows: when the interaction among the three explanatory variables was not significant (and the model had higher AIC), this interaction was removed from the model. Subsequently, the same procedure was followed for second-order interactions. GLM analyses were performed using the computer environment R (R Core Team 2017). After significance of general models, additional software (package “contrast”) was used to perform planned comparisons to address specific questions (see Results). When specific sets of data were used in multiple comparisons, their significance was corrected using the sequential Bonferroni method.

Mortality of *O. perseae* females was analysed using data from treatments containing this species. The model included the presence/absence of IG-predators, IG-prey and alternative food as explanatory variables, as well as their interactions.

IG-prey mortality was analysed using data from treatments containing IG-prey (i.e. predator juveniles). The full model included the presence/absence of the IG-predator, the herbivore and alternative food as explanatory variables, as well as their interactions.

Oviposition rates were analysed using data from treatments containing IG-predators (i.e., adult predators). The full model included the presence/absence of the herbivore, the IG prey and alternative food as explanatory variables, as well as their interactions.

## Results

### Communities with E. stipulatus as the (IG-)predator

Mortality rates of the herbivore were significantly affected by the interaction between the presence of IG-predator_ES_ and IG-prey_NC_ and between the presence of IG-prey_NC_ and pollen (Table 1a). Indeed, more prey died when IG-prey_NC_ were together with the IG-predator_ES_ than when the IG-predator_ES_ was alone (Fig 1a, compare bar 1 to 7), but not more than when the IG-prey_NC_ was alone (Fig 1a, compare bar 8 to bar 7). Also, the presence of pollen reduced herbivore mortality rates, but only in the absence of IG-prey_NC_ (Fig 1a, compare bars 4 and 6 to bars 11 and 12).

**Table 1.** Results of Generalized Linear Models applied to a) herbivore mortality rates, b) IG-prey (juveniles of *N. californicus*) mortality rates, and c) (IG-)predator (females of *E. stipulatus*) oviposition rates. All the analyses were 3 full-factorial designs. When interactions among the three explanatory variables were not significant, and if the new model yielded a lower AIC, they were removed from the model. Subsequently, the same procedure was followed for double interactions. These cases are shown in the table as NS*.

|  |  |  |  |  |  |
| --- | --- | --- | --- | --- | --- |
| a) | Herbivore mortality rates | Estimate | Std. Error | z value | Pr(>\|z\|) |
|  | Intercept | -1.755 | 0.712 | -2.466 | 0.014 |
|  | IG-predator (1) | 2.212 | 0.732 | 3.021 | 0.002 |
|  | IG-prey (2) | 2.932 | 0.729 | 4.023 | <0.001 |
|  | Pollen (3) | -1.851 | 0.609 | -3.040 | <0.001 |
|  | IG-predator * IG-prey | -2.302 | 0.756 | -3.047 | 0.002 |
|  | IG-predator * Pollen | NS |  |  |  |
|  | IG-prey * Pollen | 1.573 | 0.639 | 2.466 | .014 |
|  | (1) * (2) * (3) | NS |  |  |  |
| b) | IG-prey mortality rates | Estimate | Std. Error | z value | Pr(>\|z\|) |
|  | Intercept | 0.513 | 0.238 | 2.156 | 0.031 |
|  | IG-predator (1) | 0.591 | 0.273 | 2.163 | 0.030 |
|  | Herbivore (2) | -1.624 | 0.496 | -3.276 | 0.001 |
|  | Pollen (3) | -0.392 | 0.359 | -1.091 | 0.275 |
|  | IG-predator * Herbivore | 1.552 | 0.511 | 3.037 | 0.002 |
|  | IG-predator * Pollen | -1.705 | 0.517 | -3.300 | <0.001 |
|  | Herbivore * Pollen | 0.749 | 0.520 | 1.439 | 0.150 |
|  | (1) * (2) * (3) | NS |  |  |  |
| c) | IG-predator oviposition rates | Estimate | Std. Error | z value | Pr(>\|z\|) |
|  | Intercept | -0.843 | 0.245 | -3.443 | <0.001 |
|  | IG-prey (1) | -0.194 | 0.220 | -0.882 | 0.378 |
|  | Herbivore (2) | 0.220 | 0.216 | 1.018 | 0.308 |
|  | Pollen (3) | 1.104 | 0.235 | 4.703 | <0.001 |
|  | IG-prey * Herbivore | NS |  |  |  |
|  | IG-prey * Pollen | NS |  |  |  |
|  | Herbivore * Pollen | NS |  |  |  |
|  | (1) * (2) * (3) | NS* |  |  |  |

Mortality rates of the IG-prey_NC_ were affected by all the double interactions except that between the herbivore and pollen (Table 1b). Indeed, the presence of the IG-predator_ES_ increased the mortality of IG-prey_NC_, but only in the absence of pollen (Fig 1b, compare bars 7 and 10 to bars 11 and 13). Similarly, the presence of herbivores reduced mortality rates of IG-prey_NC_ when IG-predators_ES_ were absent (Fig1b, compare bar 8 to 9), but not when they were present (Fig1b, compare bar 7 to 10).

Planned comparisons revealed a) that IG-predators_ES_ preyed on *O. perseae* (*t*_81_ = 2.74, *P* = 0.0076; Fig 1a, compare bars 1 and 3) and marginally on the IG-prey_NC_ (*t*_80_ = −2.01, *P* = 0.048, not significant after Bonferroni correction; Fig 1b, compare bar 9 to 10) when each prey was offered alone; b) that adding IG-prey_NC_ increased mortality of *O. perseae* (*t*_81_= −2.26, *P* = 0.026; Fig 1a, compare bar 1 to 7), but adding *O. perseae* did not influence mortality of the IG-prey_NC_ (*t*_80_ = −0.31, *P* = 0.755; Fig 1b, compare bar 10 to 7); c) that the presence of pollen yielded a drastic reduction in predation of IG-predators_ES_ on both the herbivore (*t*_81_ = 2.99, *P* = 0.0037; Fig 1a, compare bar 1 to 4) and the IG-prey_NC_ (*t*_80_ = 3.91, *P* << 0.001; Fig 1b, compare bar 10 to 13); d) that when both prey were available, the presence of pollen did not affect herbivore mortality (*t*_81_ = 0.88, *P* = 0.379; Fig 1a, compare bar 7 to 11), but did lead to lower IG-prey_NC_ mortality (*t*_80_ = 3.58, *P* << 0.001; Fig 1b, compare bar 7 to 11).

Oviposition rates of IG-predators_ES_ were only affected by the presence of pollen (main factor Pollen, Table 1c). However, further planned comparisons revealed that while feeding on the herbivore stimulated egg production (*t*_96_ = 2.19, *P* = 0.021; Fig 1c, compare bar 1 to 2), feeding on IG-prey_NC_ did not (*t*_96_ = −1,13, *P* = 0.259; Fig 1c, compare bar 10 to 2).

### Communities with N. californicus as the (IG-)predator

Herbivore mortality was affected only by the interaction between IG-predator_NC_ and IG-prey_ES_ (Table 2a). Indeed, mortality of herbivores was drastically affected by the presence of IG-predators_NC_ (Fig 2a, compare bar 1 to 3), but this effect was lesser in the presence of IG-prey_ES_ (Fig 2a, compare bar 1 to 7). Mortality of IG-prey_ES_ was only affected by the presence of pollen (Table 2b).

**Table 2.** Results of Generalized Linear Models applied to a) herbivore mortality rates, b) IG-prey (juveniles of *E. stipulatus*) mortality rates, and c) (IG-)predator (females of *N. californicus*) oviposition rates. All the analyses were 3 full-factorial designs. When interactions among the three explanatory variables were not significant, and if the new model yielded a lower AIC, they were removed from the model. Subsequently, the same procedure was followed for double interactions. These cases are shown in the table as NS*.

|  |  |  |  |  |  |
| --- | --- | --- | --- | --- | --- |
| a) | Herbivore mortality rates | Estimate | Std. Error | z value | Pr(>\|z\|) |
|  | Intercept | -1.954 | 0.722 | -2.707 | 0.007 |
|  | IG-predator (1) | 2.997 | 0.729 | 4.109 | <0.001 |
|  | IG-prey (2) | 2.184 | 0.746 | 2.927 | 0.003 |
|  | Pollen (3) | -0.888 | 0.499 | -1.782 | 0.075 |
|  | IG-predator * IG-prey | -2.825 | 0.764 | -3.699 | <0.001 |
|  | IG-predator * Pollen | 0.999 | 0.460 | 2.175 | 0.030 |
|  | IG-prey * Pollen | 0.791 | 0.325 | 2.436 | 0.015 |
|  | (1) * (2) * (3) | NS* |  |  |  |
| b) | IG-prey mortality rates | Estimate | Std. Error | z value | Pr(>\|z\|) |
|  | Intercept | -0.4855 | 0.3035 | -1.600 | 0.110 |
|  | IG-predator (1) | 0.6150 | 0.3152 | 1.951 | 0.051 |
|  | Herbivore (2) | -0.3174 | 0.2851 | -1.114 | 0.265 |
|  | Pollen (3) | -1.1505 | 0.3416 | -3.368 | <0.001 |
|  | IG-predator * Herbivore | NS* |  |  |  |
|  | IG-predator * Pollen | NS* |  |  |  |
|  | Herbivore * Pollen | NS* |  |  |  |
|  | (1) * (2) * (3) | NS* |  |  |  |
| c) | IG-predator oviposition rates | Estimate | Std. Error | z value | Pr(>\|z\|) |
|  | Intercept | -2.7430 | 0.6172 | -4.444 | <0.001 |
|  | IG-prey (1) | -2.5550 | 1.0378 | -2.462 | 0.014 |
|  | Herbivore (2) | 2.5174 | 0.5989 | 4.204 | <0.001 |
|  | Pollen (3) | 0.3476 | 0.3685 | 0.943 | 0.346 |
|  | IG-prey * Herbivore | NS* |  |  |  |
|  | IG-prey * Pollen | 2.2175 | 1.1041 | 2.008 | 0.045 |
|  | Herbivore * Pollen | NS* |  |  |  |
|  | (1) * (2) * (3) | NS* |  |  |  |

Paired comparisons revealed that a) IG-predators_NC_ preyed on *O. perseae* (*t*_90_ = 3.32, *P* = 0.013; Fig 2a, compare bar 3 to 1) but not on IG-prey_ES_ (*t*_86_ = −1.35, *P* = 0.182; Fig 2b, compare bar 9 to 10), when each prey was offered alone; b) adding IG-prey_ES_ reduced mortality of *O. perseae* (*t*_90_ = 2.56, *P* = 0.012; Fig 2a, compare bar 1 to 7), but adding *O. perseae* did not change mortality of the IG-prey_ES_ (*t*_86_ = −0.93, *P* = 0.353; Fig 2b, compare bar 10 to 7); c) the presence of pollen did not affect mortality of either *O. perseae* (*t*_90_ = −0.43, *P* = 0.669; Fig 2a, compare bar 1 to 4) or the IG-prey_ES_ (*t*_86_ = 1.80, *P* = 0.075; Fig 2b, compare bar 10 to 13); d) when both types of prey were available, the presence of pollen led to a significant increase in mortality of *O.perseae* (*t*_90_ = −3.65, *P* << 0.001; Fig 2a, compare bar 7 to 11), but a significant decrease of mortality in IG-prey_ES_ (*t*_86_ = 2.04, *P* = 0.044; Fig 2b, compare bar 7 to 11).

Oviposition rates of IG-predators_NC_ were affected by the main factor Herbivore and the interaction between the IG-prey_ES_ and pollen (Table 2c). Indeed, paired comparisons revealed that e) eggs were produced when IG-predators_NC_ were offered the herbivore alone (*t*_104_ = 2.45, *P* = 0.016; Fig 2c, compare bar 1 to 2), but not when they were on arenas with either the IG-prey_ES_ (*t*_104_ = 0.01, *P* = 0.992; Fig 2c, compare bar 10 to 2) or pollen (*t*_104_ = −0.15, *P* = 0.884; Fig 2c, compare bar 5 to 2) alone. Moreover, in the presence of the herbivore, rates of oviposition were not influenced by the presence of pollen (*t*_104_ = −0.93, *P* = 0.352; Fig 2c, compare bar 1 to 4), but dramatically decreased in the presence of the IG-prey_ES_ (*t*_104_ = 2.39, *P* = 0.019; Fig 2c, compare bar 1 to 7). However, when pollen was added to the system with both prey types, IG-predators_NC_ resumed oviposition to its maximum (*t*_104_ = −2.36, *P* = 0.020; Fig 2c, compare bar 7 to 11).

## Discussion

In this study, we tested the effect of community structure on the realized interactions within a community of predatory and herbivorous mites. Because in our system the intraguild predator is the largest individual within a pair (as in most systems), we created communities in which adults (IG-predators) belonged to one species and juveniles (IG-prey) to the other, then inverted the species-stage identity in another set of communities. We then measured predation and oviposition in communities with all possible combinations of the presence of shared prey, the IG-prey, the IG-predator and an alternative food resource. We show that adding species to a community increases the number of potential trophic interactions, but not necessarily their occurrence. Indeed, despite the potential for module configurations of communities with apparent competition and intraguild predation, all modules could be described by linear food chains in our system (Box 1C).

### Basic properties of the system and implications for biocontrol

In the trophic chain configurations, although *N. californicus* killed more *O. perseae* females per day than *E. stipulatus*, oviposition rates were similar between predators. This is in line with the finding that *E. stipulatus* can only forage on mobile *O. perseae* mites when they wander outside nests, whereas *N. californicus* can penetrate inside nests and forage on all the individuals residing within (González-Fernández *et al.* 2009). This suggests that *E. stipulatus* is the most efficient predator converting food into eggs, but that *N. californicus* is more efficient at reducing herbivore populations. Which of these strategies is best for biological control will depend on the ecological condition: if outbreaks of prey are confined in time, it may be more efficient to select a biocontrol agent that feeds more, as in “inundative” biocontrol strategies, whereas controlling and keeping resident populations at low levels may be best achieved with a predator with a strong numerical response, as in “innoculative” biocontrol strategies (Van Driesche *et al.* 2007). Moreover, unlike *N. californicus, E. stipulatus* fed and oviposited on pollen. This may allow the latter to remain in the field for longer periods, as actually observed in field surveys (González-Fernández *et al.* 2009). Such temporal niche partitioning may facilitate the presence of the two predators in the same fields (Otto *et al.* 2008).

Our results also revealed asymmetry in the intraguild predation between *E. stipulatus* and *N. californicus*, with adults of the former preying upon juveniles of the latter, but not the reverse. Because *N. californicus* is likely the best competitor for the shared prey (González-Fernández *et al.* 2009), coexistence between predators is thus possible in this system (Holt & Polis 1997). Yet, the simultaneous presence of the two predators is likely to have little effect upon the densities of the shared prey. Indeed, whereas adding *N.californicus* adult intraguild predators to an arena with *E. stipulatus* juveniles results in higher shared prey densities as compared to the presence of *N. californicus* adults alone with the shared prey, the reverse is not true when adding adult *E. stipulatus* to an arena with juveniles *N. californicus*. Thus, the net effect of these interactions upon prey density is probably negligible. This is corroborated by field studies (Montserrat *et al.* 2013). However, the presence of alternative food (i.e. pollen) contributed to reduce trophic interactions between predator species resulting in community configurations that could enhance pest control. Thus, supplying alternative and preferred food to the IG-predator is probably detrimental to populations of *O. perseae*. Again, this finding is in line with field observations (Montserrat *et al.* 2013).

Optimal foraging theory predicts that species engage in trophic interactions on more than one food source when these are available (Pulliam 1974). Here, we show that *E. stipulatus* acting as intraguild predators feeds on the herbivore, *O. perseae*, on the intraguild prey, *N. californicus*, and on the alternative food, pollen, when each of these are presented alone. However, in the presence of pollen *E. stipulatus* reduces predation rates on both prey species. This may be explained by the fact that pollen is the most profitable food for this species, as found here and in other studies (Ferragut *et al.* 1987; McMurtry & Croft 1997; Bouras & Papadoulis 2005; González-Fernández *et al.* 2009). Similarly, *N. californicus* adults and juveniles ceased foraging on other food sources in presence of the herbivores. These results suggest that realized interactions hinge on the presence of the most profitable food source. In presence of the optimal food source for each of the two secondary consumers, communities tended to be reduced to two simple trophic chains. Indeed, in the most complex communities studied here, with all 5 species present, the presence of the optimal food originated the split of the community into two trophic chains, one with *E. stipulatus* feeding on pollen and the other with *N. californicus* feeding on the herbivore (Box 1 d), compare d.1.1. and d.1.2. with d.2.1. and d.2.2.).

Another factor that contributed to the linearization of the food web was that, when both the IG-prey and the shared prey were together, IG-predators_ES_ preyed mainly on the IG-prey. Indeed, mortality of *O. perseae* in presence of the IG-prey, *N. californicus*, was not affected by the presence of the IG-predator *E. stipulatus*. Furthermore, mortality of IG-prey_NC_ was significantly higher in treatments with presence of the IG-predator, compared to the control without them. This suggests that mortality in the herbivore was mainly inflicted by the IG-prey, *N. californicus*, and that the IG-predator *E. stipulatus* preyed preferentially on the IG-prey *N. californicus*. This could be explained by *E. stipulatus* having no access to *O. perseae* eggs or females located inside the nests (Montserrat *et al.* 2008a; González-Fernández *et al.* 2009), which leads to higher encounter rates between *E. stipulatus* and *N. californicus* than between *E. stipulatus* and *O. perseae*. Indeed, *E. stipulatus* forages only on mobile stages that wander outside nests (Montserrat *et al.* 2008a; González-Fernández *et al.* 2009). *Neoseiulus californicus*, however, can penetrate *O. perseae* nests, and thus may feed on them. Therefore, the realized community was that of a 4-level trophic chain (Box 1, c.2.1.). In the other community block, when *N. californicus* acted as the IG-predator, mortality of *O. perseae* females was similar in all communities with the IG-prey *E. stipulatus* present, irrespective of the presence of IG-predators_NC_. Furthermore, mortality of IG-prey_ES_ did not differ between treatments with and without the IG-predator_NC_, indicating that *N. californicus* females did not forage on *E. stipulatus* juveniles. These results suggest that, in presence of IG-prey (juveniles of *E. stipulatus*), the IG-predator_NC_ ceased to forage on either herbivore or IG-prey, likely because IG-prey_ES_ interferes with the foraging activities of IG-predators_NC_. Thus, the realized community was that of a trophic chain composed of the IG-prey, the herbivore and the plant, with the IG-predator not interacting at all (Box 1, c.2.2.). This can be explained by IG-predators_NC_ avoiding foraging on a patch where its offspring (future) IG-predator is also there. In line with this, Abad-Moyano *et al.* (2010) reported that the presence of *E. stipulatus* immatures exerted non-lethal IG-effects on *N. californicus* females, causing daily oviposition to decrease over time despite the availability of the shared prey was kept constant. In any case, here, the trophic links are again linear, with *N. californicus* being excluded from the realized community (Box 1, c.2.2.). Together, our results show that none of the complex communities was actually realized, they were all trophic chains.

### The return of the trophic chain: Fundamental vs realized trophic interactions

By combining data of mortality and oviposition at different community structures, we could determine who eats whom in a simple food web. Although this approach is powerful, it does have its limitations. Indeed, it assumes additive effects of conversion efficiencies of pairwise interactions. For example, if feeding on a prey item allows predators to better convert the food provided by another prey, this cannot be detected in our approach (i.e., indirect effects on conversion efficiency). Furthermore, it may be largely unfeasible to extend this approach to more complex food webs, although it is becoming clear that we need to know how food is transformed into predator offspring in order to fully understand food webs in nature (Neutel & Thorne 2014). Indeed, such full-factorial studies are extremely rare in the literature (but see Schmitz & Sokol-Hessner 2002; Otto *et al.* 2008).

Connectance is a fundamental measure of food-web complexity that describes the proportion of realized interactions amongst all possible ones (May 1972). It is becoming increasingly evident that connectance is generally much lower than the number of potential interactions (Beckerman *et al.* 2006). Identifying trophic links in food webs, however, is not a simple task. Molecular methods are useful to process field data and they deliver reliable information on who eats whom, but such tools currently only provide semi-quantitative estimates of predation, and they are expensive (Birkhofer *et al.* 2017). Another possible approach to measure connectance is by observations in the field (Dunne *et al.* 2002; Tylianakis *et al.* 2007; Carnicer *et al.* 2009; Lazzaro *et al.* 2009; Plein *et al.* 2013; Baiser *et al.* 2016; Lemos-Costa *et al.* 2016). Although this approach allows including a high number of species in the observations, it suffers from two main shortfalls: (a) it is generally only possible to undertake it in systems with two trophic levels in which one are primary producers, for example in plant pollinator networks (but see Bukovinszky *et al.* 2008; Neutel & Thorne 2014), or in systems where trophic interactions are detectable long after the actual events, as in parasitoid/host interactions or via the analysis of gut contents; and (b) it does not account for how foraging on a given resource translates into consumer offspring (but see Bukovinszky *et al.* 2008; Vázquez *et al.* 2015). Observations in controlled experimental settings, in contrast, deliver quantitative estimates of predation rates and concomitant offspring production, especially when trophic links, and their strength, are estimated by confronting pairs of species. Alternatively, modelling complex systems provide relative estimates on interaction strengths that go beyond pair-wise interactions (Moya-Laraño *et al.* 2012; Moya-Laraño *et al.* 2014). Yet, one-on-one approaches may ignore emergent indirect effects of having several species together (Wootton 1994; Dambacher & Ramos-Jiliberto 2007). For instance, *Cancer productus*, a crab native to the Northwest Pacific, consumes equal amounts of native oysters and of invasive drill oysters when each type of prey is offered alone, but when they are offered together crabs interact with the native oyster species only (Grason & Miner 2012). Therefore, if trophic links are not evaluated in presence of all species in the community, one may reach erroneous conclusions on the strength of the interaction (Guzmán *et al.* 2016b; Fonseca *et al.* 2017) and overestimate connectance in food webs. We show that all communities ended up becoming a sum of one or more trophic chains (Box 1C). Thus, the fundamental trophic niche of species in this system (i.e., the food items that species are potentially able to feed on) is larger than the realized trophic niche (i.e., the food items that species actually feed on when present in combinations exceeding the individual pairwise interactions (Hutchinson 1961)). Therefore, our results suggest that some food webs may be less complex than previously thought in terms of the frequency and strength of IGP.

Theoretical models exploring persistence in three-species communities with IGP find a limited parameter space for coexistence of IG-predator and IG-prey (e.g. Mylius *et al.* 2001), but field observations show that IGP is actually widespread (Polis 1991). Our results suggest that IGP in some systems might actually be occasional, as predators will tend to forage on the most profitable food, which generally is not the IG prey (Polis *et al.* 1989). In line with this, some natural systems have shown that communities with IGP actually show dynamics that are compatible with linear food chains, rather than with IGP (Borer *et al.* 2003). Therefore, predators may coexist because they rarely engage in IGP, and complexity may be over-estimated (Magalhães *et al.* 2005). This agrees with food web theory stating that weak trophic interactions promote the persistence of communities (May 1972; Paine 1992; McCann *et al.* 1998, among others). For example, Hiltunen *et al.* (2014) found long-term cycling dynamics when modelling a three-species planktonic food web with IGP, with interaction strength between IG-predator and IG-prey set to be much weaker to that between IG-predator and the shared resource. Our results suggest that increasing the number of potentially interacting species results in most species interactions becoming weaker. Indeed, the structure of interactions among species in natural communities is characterized by many weak and few strong interactions (Paine 1992; McCann *et al.* 1998), and such skewedness towards weak interactions is crucial to food web persistence (Neutel *et al.* 2002; 2007; Montoya *et al.* 2009; Neutel & Thorne 2014). Because a species’ fundamental trophic niche (all of its potential interactions) is unlikely to be realized at a particular place or time, it is crucial to determine the resources which species in a community actually feed upon, and under what circumstances. Therefore, unravelling realized food webs, (i.e., interaction strengths across different nodes and trophic levels, including indirect effects) may be key to understanding these ecological networks and their persistence.

## Acknowledgements

This preprint has been reviewed and recommended by Peer Community Ecology In (https://dx.doi.org/10.24072/pci.ecology.100008). The authors deeply thank Francis J. Burdon and two anonymous reviewers of PCI Ecology for significantly improving the quality of the manuscript with their contributions. We are indebted to Rosa María Sahún Logroño for her valuable help in maintaining the experimental populations. This work was financed by the Spanish Ministry of Science and Innovation (CGL2015-66192-R and AGL2011-30538-C03-03). I.T.C. was recipient of a grant from the Spanish National Research Council (CSIC, ref: JAE-PRE 041). The animals used for the research of this publication are not test animals in the legal sense. The authors have no conflict of interest to declare.

## Conflict of interest disclosure

The authors of this preprint declare that they have no financial conflict of interest with the content of this article. Sara Magalhães, Jordi Moya-Laraño and Marta Montserrat are recommenders at PCI Ecology.

---

**Box 1.**
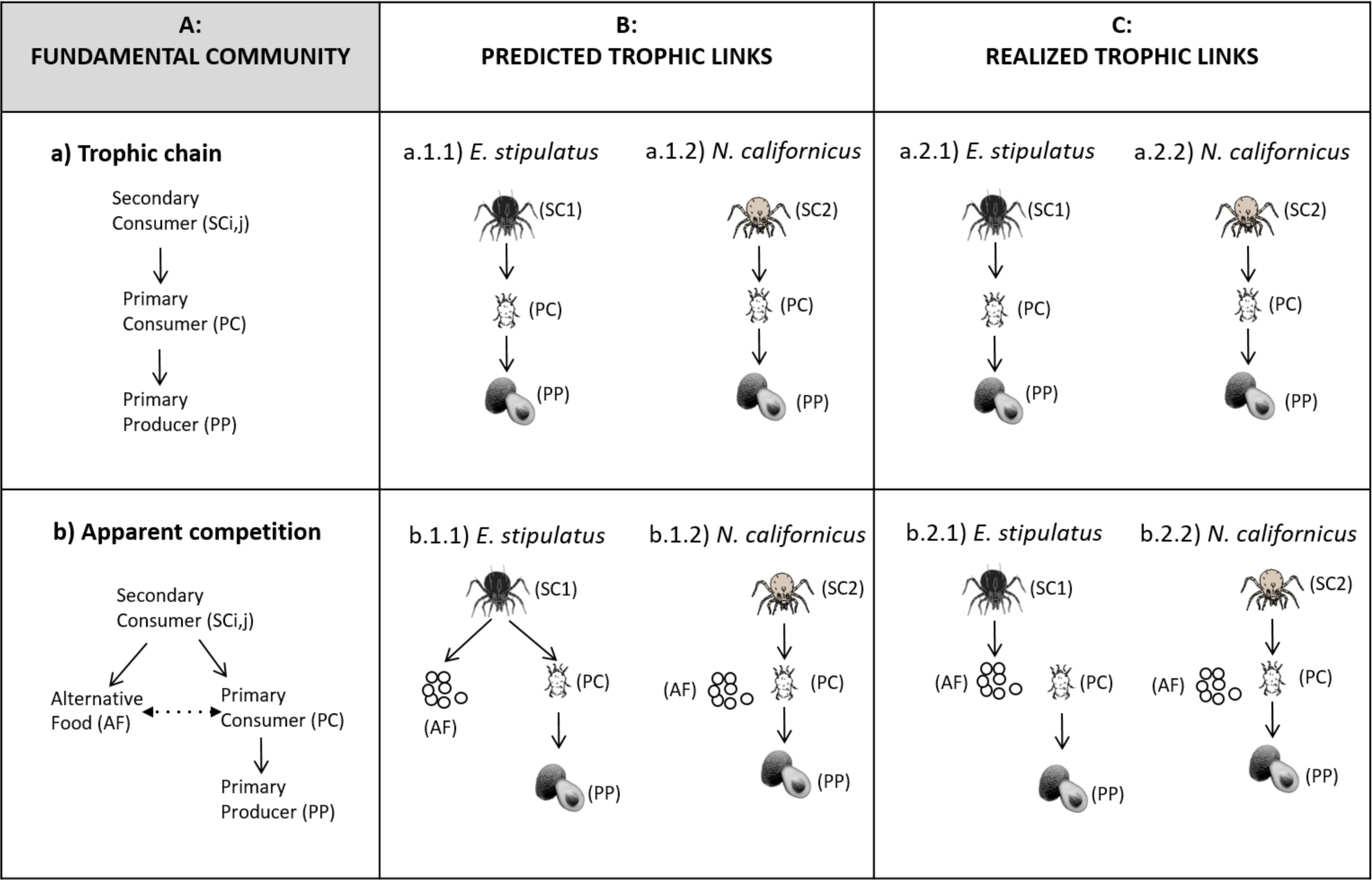
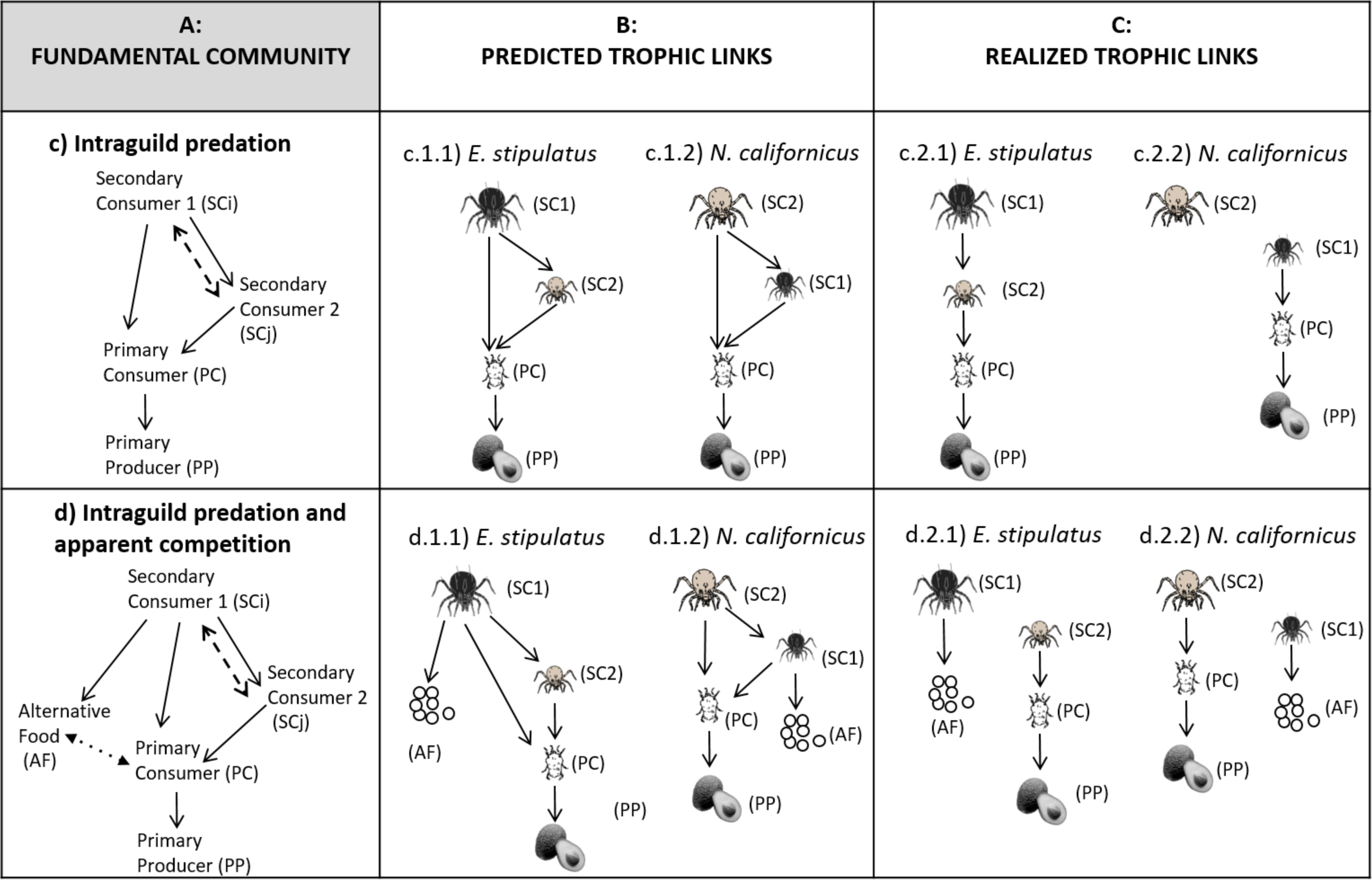
**A: Fundamental community modules** included in this study. a) trophic chain, b) apparent competition, c) intraguild predation, and d) intraguild predation and apparent competition. From a) to d) the complexity of the community is increased via increasing the number of species and the number of interactions among them. **B: Predicted trophic links** that have been observed using pairwise experimental settings**. C: Realized trophic links** occurring across community modules of increasing complexity, obtained from the experiments presented here, where interactions are measured in the presence of other components of the community. SC stands for secondary consumer, PC for primary consumer, PP for primary producer, and AF for alternative food. SC1 and SC2 are phytoseiid predatory mites, i.e. *Euseius stipulatus* and *Neoseiulus californicus*, respectively, PC is the tetranychid herbivore mite *Oligonychus perseae*, AF is pollen of *Carpobrotus edulis*, and PP is the avocado *Persea americana*. Solid arrows indicate negative direct interactions (who eats whom), whereas dotted and dashed arrows in Box 1A indicate negative indirect interactions (apparent competition and competition).

## References

Abad-Moyano, R., Urbaneja, A. & Schausberger, P. (2010). Intraguild interactions between Euseius stipulatus and the candidate biocontrol agents of Tetranychus urticae in Spanish clementine orchards: Phytoseiulus persimilis and Neoseiulus californicus. Experimental and Applied Acarology, 50, 23–34.

Amarasekare, P. (2008). Coexistence of intraguild predators and prey in resource-rich environments. Ecology, 89, 2786–2797.

Baiser, B., Elhesha, R. & Kahveci, T. (2016). Motifs in the assembly of food web networks. Oikos, 125, 480–491.

Bascompte, J. & Melián, C.J. (2005). Simple trophic modules for complex food webs. Ecology, 86, 2868–2873.

Bascompte, J., Jordano, P. & Olesen, J.M. 2006) Asymmetric co-evolutionary networks facilitate biodiversity maintenance. Science, 312, 431–433.

Beckerman, A.P., Petchey, O.L. & Warren, P.H. (2006). Foraging biology predicts food web complexity. Proceedings of the National Academy of Sciences, 103, 13745–13749.

Birkhofer, K., Bylund, H., Dalin, P., Ferlian, O., Gagic, V., Hambäck, P.A. et al. (2017). Methods to identify the prey of invertebrate predators in terrestrial field studies. Ecology and Evolution, 7, 1942–1953.

Borer, E.T., Briggs, C.J., Murdoch, W.W. & Swarbrick, S.L. (2003). Testing intraguild predation theory in a field system: does numerical dominance shift along a gradient of productivity? Ecology Letters, 6, 929–935.

Bouras, S.L. & Papadoulis, G.T. (2005). Influence of selected fruit tree pollen on life history of Euseius stipulatus (Acari: Phytoseiidae). Experimental and Applied Acarology, 36, 1–14.

Bukovinszky, T., van Veen, F.F., Jongema, Y. & Dicke, M. (2008). Direct and indirect effects of resource quality on food web structure. Science, 319, 804–807.

Carnicer, J., Jordano, P. & Melián, C.J. (2009). The temporal dynamics of resource use by frugivorous birds: a network approach. Ecology, 90, 1958–1970.

Carpenter, S.R., Kitchell, J.F. & Hodgson, J.R. (1985). Cascading trophic interactions and lake productivity. BioScience, 35, 634–639.

Dambacher, J.M. & Ramos-Jiliberto, R. (2007). Understanding and predicting effects of modified interactions through a qualitative analysis of community structure. The Quarterly Review of Biology, 82, 227–250.

Daugherty, M.P., Harmon, J.P. & Briggs, C.J. (2007). Trophic supplements to intraguild predation. Oikos, 116, 662–677.

Diehl, S. & Feißel, M. (2000). Effects of enrichment on three-level food chains with omnivory. The American Naturalist, 155, 200–218.

Dunne, J.A., Williams, R.J. & Martinez, N.D. (2002). Food-web structure and network theory: the role of connectance and size. Proceedings of the National Academy of Sciences, 99, 12917–12922.

Elton, C. (1927) Animal Ecology. Sidgwick and Jackson, London. The University of Chicago Press, ISBN 0-226-20639-4

Emmerson, M. & Yearsley, J.M. (2004). Weak interactions, omnivory and emergent food-web properties. Proceedings of the Royal Society of London B: Biological Sciences, 271, 397–405.

Faria, L.D.B. & Costa, M.I.S. (2010). Omnivorous food web, prey preference and allochthonous nutrient input. Ecological Complexity, 7, 107–114.

Ferragut, F., Garcia-Mari, F., Costa-Comelles, J. & Laborda, R. (1987). Influence of food and temperature on development and oviposition of Euseius stipulatus and Typhlodromus phialatus (Acari: Phytoseiidae). Experimental and Applied Acarology, 3, 317–329.

Fonseca, M.M., Montserrat, M., Guzmán, C., Torres-Campos, I., Pallini, A. & Janssen, A. (2017). How to evaluate the potential occurrence of intraguild predation. Experimental and Applied Acarology, 72, 103–114.

Gagnon, A.E., Heimpel, G.E. & Brodeur, J. (2011). The ubiquity of intraguild predation among predatory arthropods. PLoS ONE, 6, e28061.

Gellner, G. & McCann, K. (2012). Reconciling the omnivory-stability debate. The American Naturalist, 179, 22–37.

Gellner, G. & McCann, K.S. (2016). Consistent role of weak and strong interactions in high-and low-diversity trophic food webs. Nature communications, 7, 11180.

González-Fernández, J., De la Peña, F., Hormaza, J.I., Boyero, J., Vela, J., Wong, E. et al. (2009). Alternative food improves the combined effect of an omnivore and a predator on biological pest control. A case study in avocado orchards. Bulletin of Entomological Research, 99, 433– 444.

Grason, E.W. & Miner, B.G. (2012). Preference alters consumptive effects of predators: top-down effects of a native crab on a system of native and introduced prey. PloS ONE, 7, e51322.

Guzmán, C., Aguilar-Fenollosa, E., Sahún, R.M., Boyero, J.R., Vela, J.M., Wong, E. et al. (2016a). Temperature-specific competition in predatory mites: Implications for biological pest control in a changing climate. Agriculture, Ecosystems and Environment, 216, 89–97.

Guzmán, C., Sahún, R. & Montserrat, M. (2016b). Intraguild predation between phytoseiid mite species might not be so common. Experimental and Applied Acarology, 68, 441–453.

Hairston, N.G., Smith, F.E. & Slobodkin, L.B. (1960). Community structure, population control, and competition. The American Naturalist, 94, 421–425.

Heithaus, M.R. (2001). Habitat selection by predators and prey in communities with asymmetrical intraguild predation. Oikos, 92, 542–554.

Hiltunen, T., Ellner, S.P., Hooker, G., Jones, L.E. & Hairston Jr, N.G. (2014). Eco-evolutionary dynamics in a three-species food web with intraguild predation: intriguingly complex. In: Advances in Ecological Research. Elsevier, pp. 41–73.

Hin, V., Schellekens, T., Persson, L. & de Roos, A.M. (2011). Coexistence of predator and prey in intraguild predation systems with ontogenetic niche shifts. The American Naturalist, 178, 701–714.

Holt, R.D. (1977). Predation, apparent competition, and the structure of prey communities. Theoretical Population Biology, 12, 197–229.

Holt, R.D. (1997). Community modules. In: Multitrophic interactions in Terrestrial Systems (eds. Gange, A & Brown, V). Blackwell Science London, UK, pp. 333–350.

Holt, R.D. & Huxel, G.R. (2007). Alternative prey and the dynamics of intraguild predation: theoretical perspectives. Ecology, 88, 2706–2712.

Holt, R.D. & Polis, G.A. (1997). A theoretical framework for intraguild predation. The American Naturalist, 149, 745–764.

Hutchinson, G. E. 1957. Concluding remarks. Cold Spring Harbor Symposia on Quantitative Biology 22: 415–427.

Hutchinson, G.E. (1961). The paradox of the plankton. The American Naturalist, 95, 137–145.

Janssen, A., Montserrat, M., HilleRisLambers, R., Roos, A.M., Pallini, A. & Sabelis, M.W. (2006). Intraguild predation usually does not disrupt biological control. In: Trophic and Guild in Biological Interactions Control (eds. Brodeur, J & Boivin, G). Springer Netherlands Dordrecht, pp. 21–44.

Janssen, A., Sabelis, M.W., Magalhaes, S., Montserrat, M. & van der Hammen, T. (2007). Habitat structure affects intraguild predation. Ecology, 88, 2713–2719.

Johnson, S., Domínguez-García, V., Donetti, L. & Muñoz, M. A. Trophic coherence determines foodweb stability. Proc.Natl. Acad. Sci. 111, 17923–17928.

Kondoh, M. (2008). Building trophic modules into a persistent food web. Proceedings of the National Academy of Sciences, 105, 16631–16635.

Kratina, P., Hammill, E. & Anholt, B.R. (2010). Stronger inducible defences enhance persistence of intraguild prey. Journal of Animal Ecology, 79, 993–999.

Křivan, V. (2000). Optimal intraguild foraging and population stability. Theoretical Population Biology, 58, 79–94.

Křivan, V. & Diehl, S. (2005). Adaptive omnivory and species coexistence in tri-trophic food webs. Theoretical Population Biology, 67, 85–99.

Kuijper, L.D., Kooi, B.W., Zonneveld, C. & Kooijman, S.A. (2003). Omnivory and food web dynamics. Ecological Modelling, 163, 19–32.

Lazzaro, X., Lacroix, G., Gauzens, B., Gignoux, J. & Legendre, S. (2009). Predator foraging behaviour drives food-web topological structure. Journal of Animal Ecology, 78, 1307–1317.

Lemos-Costa, P., Pires, M.M., Araújo, M.S., Aguiar, M.A. & Guimarães, P.R. (2016). Network analyses support the role of prey preferences in shaping resource use patterns within five animal populations. Oikos, 125, 492–501.

Liu, Z. & Zhang, F. (2013). Species coexistence of communities with intraguild predation: The role of refuges used by the resource and the intraguild prey. Biosystems, 114, 25–30.

Magalhães, S., Tudorache, C., Montserrat, M., Van Maanen, R., Sabelis, M.W. & Janssen, A. (2005). Diet of intraguild predators affects antipredator behavior in intraguild prey. Behavioral Ecology, 16, 364–370.

May, R.M. (1972). Will a large complex system be stable? Nature, 238, 413.

McCann, K., Hastings, A. & Huxel, G.R. (1998). Weak trophic interactions and the balance of nature. Nature, 395, 794.

McMurtry, J. & Croft, B. (1997). Life-styles of phytoseiid mites and their roles in biological control. Annual Review of Entomology, 42, 291–321.

Messelink, G.J. & Janssen, A. (2014). Increased control of thrips and aphids in greenhouses with two species of generalist predatory bugs involved in intraguild predation. Biological Control, 79, 1–7.

Montoya, J., Woodward, G., Emmerson, M.C. & Solé, R.V. (2009). Press perturbations and indirect effects in real food webs. Ecology, 90, 2426–2433.

Montserrat, M., de la Peña, F., Hormaza, J. & González-Fernández, J. (2008a). How do Neoseiulus californicus (Acari: Phytoseiidae) females penetrate densely webbed spider mite nests? Experimental and Applied Acarology, 44, 101–106.

Montserrat, M., Guzmán, C., Sahún, R., Belda, J. & Hormaza, J. (2013). Pollen supply promotes, but high temperatures demote, predatory mite abundance in avocado orchards. Agriculture, Ecosystems & Environment, 164, 155–161.

Montserrat, M., Magalhães, S., Sabelis, M.W., De Roos, A.M. & Janssen, A. (2008b). Patterns of exclusion in an intraguild predator–prey system depend on initial conditions. Journal of Animal Ecology, 77, 624–630.

Moya-Laraño, J., Bilbao-Castro, J.R., Barrionuevo, G., Ruiz-Lupión, D., Casado, L.G., Montserrat, M. et al. (2014). Eco-evolutionary spatial dynamics: rapid evolution and isolation explain food web persistence. Advances in Ecological Research, 50, 75–144.

Moya-Laraño, J., Verdeny-Vilalta, O., Rowntree, J., Melguizo-Ruiz, N., Montserrat, M. & Laiolo, P. (2012). Climate Change and Eco-Evolutionary Dynamics in Food Webs. Advances in Ecological Research, 47, 1–80.

Mylius, S.D., Klumpers, K., de Roos, A.M. & Persson, L. (2001). Impact of intraguild predation and stage structure on simple communities along a productivity gradient. The American Naturalist, 158, 259–276.

Nakazawa, T., Miki, T. & Namba, T. (2010). Influence of predator-specific defense adaptation on intraguild predation. Oikos, 119, 418–427.

Neutel, A.M., Heesterbeek, J.A. & de Ruiter, P.C. (2002). Stability in real food webs: weak links in long loops. Science, 296, 1120–1123.

Neutel, A.M., Heesterbeek, J.A., Van de Koppel, J., Hoenderboom, G., Vos, A., Kaldeway, C. et al. (2007). Reconciling complexity with stability in naturally assembling food webs. Nature, 449, 599.

Neutel, A.M. & Thorne, M.A. (2014). Interaction strengths in balanced carbon cycles and the absence of a relation between ecosystem complexity and stability. Ecology letters, 17, 651–661.

Novak, M. (2013). Trophic omnivory across a productivity gradient: intraguild predation theory and the structure and strength of species interactions. Proceedings of the Royal Society of London B: Biological Sciences, 280, 20131415.

Novak, M. & Wootton, J.T. (2010). Using experimental indices to quantify the strength of species interactions. Oikos, 119, 1057–1063.

Oksanen, L., Fretwell, S.D., Arruda, J. & Niemela, P. (1981). Exploitation ecosystems in gradients of primary productivity. The American Naturalist, 118:2, 240-261.

Otto, S.B., Berlow, E.L., Rank, N.E., Smiley, J. & Brose, U. (2008). Predator diversity and identity drive interaction strength and trophic cascades in a food web. Ecology, 89, 134–144.

Paine, R.T. (1992). Food-web analysis through field measurement of per capita interaction strength. Nature, 355, 73.

Pimm, S. L. (1984). Complexity and stability of ecosytems. Nature, 307, 321–326.

Pimm, S.L. & Lawton, J. (1978). On feeding on more than one trophic level. Nature, 275, 542–544.

Plein, M., Längsfeld, L., Neuschulz, E.L., Schultheiß, C., Ingmann, L., Töpfer, T. et al. (2013). Constant properties of plant–frugivore networks despite fluctuations in fruit and bird communities in space and time. Ecology, 94, 1296–1306.

Polis, G.A. (1991). Complex trophic interactions in deserts: an empirical critique of food-web theory. The American naturalist, 138, 123–155.

Polis, G.A. & Holt, R.D. (1992). Intraguild predation: the dynamics of complex trophic interactions. Trends in Ecology & Evolution, 7, 151–154.

Polis, G.A., Myers, C.A. & Holt, R.D. (1989). The ecology and evolution of intraguild predation: potential competitors that eat each other. Annual Review of Ecology and Systematics, 20, 297–330.

Polis, G.A. & Strong, D.R. (1996). Food web complexity and community dynamics. The American Naturalist, 147, 813–846.

Prill R.J., Iglesias P.A., Levchenko A. (2005). Dynamic properties of network motifs contribute to network organization. PLOS BIOLOGY, 3: 1881-1892.

Pulliam H.R. (1974). On the thory of optimal diets. American Naturalist, 108: 59-74.

R Core Team (2017). R: A Language and Environment for Statistical Computing. https://www.R-project.org/

Rosenheim, J.A., Kaya, H., Ehler, L., Marois, J.J. & Jaffee, B. (1995). Intraguild predation among biological-control agents: theory and evidence. Biological Control, 5, 303–335.

Rudolf, V.H. (2007). The interaction of cannibalism and omnivory: consequences for community dynamics. Ecology, 88, 2697–2705.

Rudolf, V.H.W. & Armstrong, J. (2008). Emergent impacts of cannibalism and size refuges in prey on intraguild predation systems. Oecologia, 157, 675–686.

Schmitz, O.J. & Sokol-Hessner, L. (2002). Linearity in the aggregate effects of multiple predators in a food web. Ecology Letters, 5, 168–172.

Tylianakis, J.M., Tscharntke, T. & Lewis, O.T. (2007). Habitat modification alters the structure of tropical host–parasitoid food webs. Nature, 445, 202.

Van Driesche, R., Hoddle, M. & Center, T.D. (2007). Control de plagas y malezas por enemigas naturales. US Department of Agriculture, US Forest Service, Forest Health Technology Enterprise Team.

Vance-Chalcraft, H.D., Rosenheim, J.A., Vonesh, J.R., Osenberg, C.W. & Sih, A. (2007). The influence of intraguild predation on prey suppression and prey release: a meta-analysis. Ecology, 88, 2689–2696.

Vázquez, D.P., Ramos-Jiliberto, R., Urbani, P. & Valdovinos, F.S. (2015). A conceptual framework for studying the strength of plant–animal mutualistic interactions. Ecology letters, 18, 385–400.

Wootton, J.T. (1994). Putting the pieces together: testing the independence of interactions among organisms. Ecology, 75, 1544–1551.

Wootton, J.T. & Emmerson, M. (2005). Measurement of interaction strength in nature. Annual Review of Ecology, Evolution, and Systematics, 36, 419–444.

